# Elevated estradiol during a hormone simulated pseudopregnancy decreases sleep and increases hypothalamic activation in female Syrian hamsters

**DOI:** 10.1101/2022.10.27.514063

**Authors:** Abiola Irvine, Maeve I. Gaffney, Erin K. Haughee, Marité A. Horton, Hailey C. Morris, Kagan C. Harris, Jaclyn E. Corbin, Clara Merrill, Michael L. Perlis, Laura E. Been

## Abstract

Sleep disruptions are a common occurrence during the peripartum period. While physical and environmental factors associated with pregnancy and newborn care account for some sleep disruptions, there is evidence that peripartum fluctuations in estrogens may independently impact sleep. We therefore used a hormone-simulated pseudopregnancy in female Syrian hamsters to test the hypothesis that pregnancy-like increases in estradiol decrease sleep in the absence of other factors. Adult female Syrian hamsters were ovariectomized and given daily hormone injections that simulate estradiol levels during early pregnancy, late pregnancy, and the postpartum period. Home cage video recordings were captured at seven timepoints and videos were analyzed for actigraphy. During “late pregnancy,” total sleep time and sleep efficiency were decreased in hormone-treated animals during the white light period compared to vehicle controls. During both “early pregnancy” and “late pregnancy,” locomotion was increased in the white light period for hormone-treated animals; this change continued into the “postpartum period” for animals who continued to receive estradiol treatment, but not for animals who were withdrawn from estradiol. At the conclusion of the experiment, animals were euthanized and cFos expression was quantified in the ventral lateral preoptic area (VLPO) and lateral hypothalamus (LH). Animals who continued to receive high levels of estradiol during the “postpartum” period had significantly more cFos in the VLPO and LH than animals who were withdrawn from hormones or vehicle controls. Together, these data suggest that increased levels of estradiol during pregnancy are associated with sleep suppression which may be mediated by increased activation of hypothalamic nuclei.

## Introduction

Sleep complaints are a common, but potentially serious, occurrence during pregnancy and the postpartum period. Nearly all pregnant people report experiencing nighttime awakenings during pregnancy^1^, while up to three quarters report poor sleep quality during pregnancy^2–4^, and more than half report symptoms of insomnia^1,5^. As pregnancy progresses, sleep disturbances increase, such that there are more sleep disturbances in the third trimester compared with the first two trimesters^6–8^. These disruptions often reach clinically-significant levels; the prevalence of diagnosed insomnia, sleep-disordered breathing, and restless leg syndrome (RLS) are all elevated during pregnancy^9,10^. Poor sleep quality in late pregnancy is predictive of poor sleep quality in the postpartum period^11^, and sleep disturbances during pregnancy are associated with adverse maternal and fetal outcomes^12^, including heightened risk for gestational diabetes^13^, increased pain during labor^14^, more cesarean sections^15^, preterm birth^16^, low birth weight^17^, and more depressive symptoms during the postpartum period^11,18,19^. Despite these serious consequences, the pathophysiology of peripartum sleep disorders is poorly understood.

It is easy to assume that sleep disruptions during the peripartum period are due solely to the dramatic anatomical, physiological, and environmental changes that accompany pregnancy and new parenthood. Indeed, increased body weight and uterine volume during pregnancy make it more difficult to breathe comfortably when lying down, and changes in body shape may necessitate more frequent repositioning, affecting sleep continuity^20^. Pregnancy symptoms such as nausea, heartburn, round ligament pain, and increased urinary frequency also contribute to disruptions in sleep during pregnancy^8^. Following delivery, the demands of newborn care significantly disrupt sleep, especially during the first month postpartum^21^. However, there is evidence that in addition to these anatomical and environmental factors, peripartum hormone fluctuations may influence sleep through direct actions on the brain.

Several hormones including estrogens, progesterone, oxytocin, endorphins, and corticosterones all fluctuate during the peripartum period^22^. Among these hormones, estrogens stand out as particularly strong candidates to influence peripartum sleep. First, there are known sex differences in sleep in rodents and humans^23,24^. In rodents, these sex differences are eliminated by ovariectomy^25,26^, and acute exogenous administration of estradiol decreases total sleep time and REM Sleep^27,28^, suggesting ovarian hormones as a primary mediator of sleep in females. In support of this, sleep varies across the menstrual cycle in humans and estrous cycle in rodents^29^. Notably, in both humans and rodents, sleep is most disturbed at the points in the cycle when estradiol levels are elevated^30^, suggesting that high levels of estrogens may inhibit sleep. Compared with fluctuations in estrogens across the menstrual or estrous cycle, peripartum fluctuations in estrogens are much larger; in humans, estrogen levels rise precipitously across pregnancy, peaking during the third trimester at 100-1000-fold higher than pre-pregnancy levels^31^. Following birth and the expulsion of the placenta, however, estrogen levels quickly drop to below pre-gravid levels and remain suppressed until ovulation resumes^32^. However, the impact of these large fluctuations in estrogens on peripartum sleep is unclear, primarily because it is difficult to tease apart the direct effects of estrogens on sleep from those resulting from the growth and development of the fetus or associated with parental care^23^.

In the brain, estrogen receptors (ERs) are localized in many sleep/wake-regulatory regions, including several hypothalamic nuclei, the dorsal raphe, and the locus coeruleus^33,34^. Hypothalamic sleep-wake regulatory nuclei such as the ventral lateral preoptic nucleus (VLPO) and lateral hypothalamus (LH) are strong putative candidates to mediate peripartum sleep disruptions. First, the VLPO and LH have dense concentrations of ERs^33,34^, suggesting high sensitivity to fluctuations in estrogens. Estradiol inhibits activation of sleep-promoting VLPO neurons^35^ and reduces the expression of sleep-promoting signaling molecules in the VLPO^36–38^. In contrast, sleep deprivation increases cFos expression in the LH, and this increase is modulated by estradiol administration in ovariectomized female rats^39^. The independent impact of peripartum estradiol fluctuations on the activation of sleep nuclei, however, has not been investigated. We therefore hypothesized that high levels of estradiol during pregnancy would decrease sleep measures via decreased VLPO activity and increased LH activity. To test this hypothesis, we used a hormone simulated pseudopregnancy (HSP) in female Syrian hamsters to directly test the impact of peripartum estradiol fluctuations on sleep actigraphy measures and cFos expression in the VLPO and LH.

## Materials and Methods

### Subjects

Adult female Syrian hamsters (*Mesocricetus auratus*, n = 24) were ordered from Charles Rivers Laboratories (Wilmington, MA, USA) at approximately 60 days old. All hamsters were housed individually in polycarbonate cages (19” × 10.5” × 8”) with aspen bedding. Hamsters are solitary, territorial animals and individual housing is standard and non-stressful^40,41^. Animal holding rooms were temperature- and humidity-controlled and maintained on a reversed 14-hr light/10-hr dark period, which is a common housing condition for Syrian hamsters and simulates the longer “summer-like” days during which female hamsters would breed in the wild^42^. Food and water were provided *ad libitum*. All animal procedures were carried out in accordance with the National Institutes of Health Guide for the Care and Use of Laboratory Animals and approved by the Institutional Animal Care and Use Committee.

### Ovariectomy

In order to remove the endogenous source of circulating estrogen and progesterone, females were bilaterally ovariectomized prior to the initiation of the hormone simulated pseudopregnancy (see below). Surgery was conducted using aseptic surgical technique and under isoflurane anesthesia (2-3% vaporized in oxygen, Piramal, Bethlehem, PA, USA). Analgesic (Meloxicam, 2mg/kg, Portland, ME, USA) was administered subcutaneously immediately prior to the start of surgery and for three days postoperatively. Briefly, subjects’ bilateral flanks were shaved and cleaned with three alternating scrubs of 70% ethanol and betadyne before being transferred to a sterile surgical field. Anesthesia was maintained via a nosecone. Subjects’ ovaries were extracted via bilateral flank incisions and removed via cauterization of the uterine horn and blood vessels. Polydioxanone absorbable suture (Ethicon, Sommerville, NJ, USA) and wound clips (Fine Science Tools, Foster City, CA, USA) were used to close the smooth muscle and skin incisions, respectively.

### Hormone Simulated Pseudopregnancy

Following recovery from ovariectomy (see Surgery), the HSP (HSP) protocol was initiated. In this model, ovariectomized females are given daily injections of estrogen and/or progesterone. The timing and doses were initially chosen for their ability to induce maternal behaviors in nulliparous ovariectomized rats^43–45^, but have since been used to model peripartum fluctuations in ovarian hormones ^46–51^ and modified to fit the gestational profile of other rodent species^52,53^ including Syrian hamsters^54^. All hamsters were administered daily subcutaneous injections of hormone dissolved in cottonseed oil vehicle (Sigma, St. Louis, MO, USA) over a 22-day period. On days 1-12 (“early pregnancy”), hamsters received a low dose (2.5 μg) of estradiol benzoate (Sigma) and a high dose (4 mg) of progesterone (Sigma). On days 13-17 (“late pregnancy”), hamsters received a high dose of estradiol benzoate (50 μg). On days 18-22, hamsters were divided into two experimental conditions: estrogen-withdrawn (“withdrawn” n = 8) females received daily vehicle injections, modeling the dramatic drop in estrogen that occurs during the postpartum period, while females in the estrogen-sustained (“sustained” n = 8) group continued to receive daily injections of a high dose (50 μg) of estradiol. Finally, both withdrawn and sustained females were compared to a “no hormone” condition (n = 8), in which females received daily vehicle (oil) injections for all 22 days of the hormone simulated pregnancy.

### Actigraphy

Behavioral actigraphy measures were used as a proxy for sleep. Subjects were video recorded in their home cages for twenty-four-hour blocks using Logitech C922 Prostream Web Cameras (Logitech, Lausanne, Switzerland) streaming to Acer Travelmate Pro computers (Acer, Hsinchu, Taiwan) running OBS Studio software (Open Broadcaster Studio, https://obsproject.com). Recordings were collected at seven different times corresponding to the HSP protocol: 2 pre-test recordings prior to the initiation of HSP (Days -2 and -1); 2 recordings during “early pregnancy” (EP, Days 5 and 10); 2 recordings during “late pregnancy” (LP, Days 16 and 17), and one recording during the “postpartum period” (Day 22). Data files were saved to Box cloud-based storage (Box Inc., Redwood City, CA, USA) after each recording session ended, and then downloaded to a local computer with EthoVision XT software (version 14, Noldus Information Technology, Wageningen, The Netherlands) for actigraphy analyses.

Ethovision uses grayscale pixel values to differentiate the subject from the background and subsequently calculate activity states. Each pixel was assigned a grayscale value using a scale of Black = 0 and White = 200. A reference image from each video was used to assign grayscale values to the hamster, and the “dynamic subtraction” feature was used to differentiate the hamster (darker) from the background (lighter). Activity was defined using the following formula: Activity = (CP_n_/P_n_) × 100, where CP = Changed Pixels and P = Total Pixels. As such, activity is equal to the percentage change in all pixels. An activity value was calculated for each frame, allowing for tracking of activity state across time. Arena settings were created in Ethovision for each cage individually to exclude the food hopper and any reflective metal from the analysis arena.

Several filters were applied to smooth the data and increase the accuracy of activity state detection. Data profile filters were applied to adjust the following parameters: sample number (the number of samples used to calculate the moving average), threshold percentage (the percent change threshold used for pixel comparison); and seconds (the number of seconds a state must last before it is classified as active or inactive); if the duration did not exceed the threshold, the previous state (active or inactive) ended, but no new state was initiated. Separate data profile filters were applied to the light phase and dark phase to optimize accurate tracking under different illumination conditions. In the light phase (white illumination), the following filters were applied: 15 samples, 0.70% threshold, 80 seconds. In the dark phase (red illumination), the following filters were applied: 30 samples, 0.03% threshold, 150 seconds.

Under both light conditions, the activity threshold was set to 5, meaning that there needed to be a 5-pixel difference in grayscale value for an animal to be classified as active. The background noise filter was set to 1, meaning a change in pixel was only counted if the adjacent pixel also changed. Finally, movement filters were applied to all videos. The state was considered “moving” if the running average velocity exceeded 0.80 cm/s (start velocity) and would remain classified as “moving” until the running average velocity dropped below 0.05 cm/s (stop velocity). Instances of movement shorter than 0.10 s were excluded and 6 samples were averaged to determine movement state.

Using these parameters, the following dependent variables were collected: total sleep time (cumulative seconds inactive), total number of sleep bouts (frequency of inactive state), and distance traveled (cumulative cm moved from center point). In addition, sleep efficiency (cumulative duration of inactivity /total sleep period) and wake after sleep onset (WASO; latency to achieve active state during the white light period) were calculated. Across data collection, six video recordings were lost due to technical failure and were excluded from actigraphy analyses.

### Histology and Immunohistochemistry

#### Perfusion

On Day 23, subjects were sacrificed by intracardial perfusion. Subjects were given an overdose of sodium pentobarbital (Beuthanasia-D Special, 22 mg/100g body weight, Merck Animal Health, Madison, NJ, USA). Once at surgical plane, subjects were transcardially perfused with approximately 200 mL of 25 mM PBS (pH 7.4), followed by 200 mL of 4% paraformaldehyde (Electron Microscopy Sciences, Hatfield, PA, USA). Brains were immediately removed and post-fixed in 4% paraformaldehyde overnight (4° C) and then cryoprotected for 48 hr in 30% sucrose in PBS. Coronal sections (35-μm) of brain tissue were sectioned on a cryostat (−20°C), collected in a 1:4 series, and stored in cryoprotectant until immunohistochemical processing.

#### Immunohistochemistry

Sections were removed from cryoprotectant and rinsed 5 × 5 min in 25 mM PBS. To reduce endogenous peroxidase activity, tissue sections were incubated in 0.3% hydrogen peroxide for 15 min. After 5 × 5 min rinses in PBS, sections were incubated in a rabbit monoclonal primary antibody against cFos (1:10,000, 9F6 Rabbit mAb #2250, Cell Signaling, Danvers, MA, USA) in 0.4% Triton-X 100 for 48 hr at room temperature. After incubation in primary antibody, sections were rinsed in PBS and then incubated for 1 hr in a biotinylated secondary antibody (anti-rabbit, 1:600 Jackson ImmunoResearch, West Grove, PA, USA) in PBS with 0.4% Triton-X 100. Sections were rinsed again in PBS and then incubated for 1 hr in avidin-biotin complex (4.5 μL each of A and B reagents/ml PBS with 0.4% Triton-X 100, ABC Elite Kit, Vector Laboratories, Burlingame, CA, USA). After rinsing in PBS, sections were incubated in 3,3’-diaminobenzidine HCl (0.2 mg/mL, Sigma) and hydrogen peroxide (0.83 μL/mL, Sigma) for 10 min, yielding an orange-brown product. The reaction was stopped by rinsing sections in PBS. A subset of sections was run through the immunohistochemistry protocol with the primary antibody omitted. Stained tissue sections were mounted onto subbed glass slides and allowed to air-dry overnight. Slides were then dehydrated in alcohols, cleared in xylenes, and cover slipped using Permount (Fisher Scientific, Waltham, MA, USA).

#### Microscopy and Cell Counts

A light microscope (Eclipse E200, Nikon, Tokyo, Japan) using a color camera and SpotBasic software (Diagnostic Instruments, Sterling Heights, MI, USA) was used to acquire images. Using the Stereotaxic Atlas of The Golden Hamster Brain^55^ as a reference and the anterior commissure (ac), optic chiasm (ox)/optic tract (ot), third ventricle (3V), and shape of the cortex were used as neuroanatomical landmarks (**Figure 5**). 10X images of the VLPO and LH were acquired for each animal at two levels: rostral VLPO (atlas figure 19-20), caudal VLPO (atlas figure 21-22), rostral LH (atlas figure 26-27), and caudal LH (atlas figure 31-32). Prior to counting, images were converted to grayscale and enhanced for contrast. cFos-immunoreactive (-ir) cells were counted manually by experimenters blind to the condition of the subject and inter-rater reliability was confirmed.

### Statistical analysis

Data from the light versus dark condition were analyzed separately. Repeated Measures 2-Way ANOVAs were used to examine the interaction of hormone condition (oil, estrogen-sustained, estrogen-withdrawn) and HSP day on each of the dependent variables (cumulative sleep duration, sleep efficiency, number of sleep bouts, WASO, and distance moved). Tukey’s HSD post-hoc tests were used to explicate significant effects. One-way ANOVAs were used to determine potential effects of hormonal condition on the number of cFos-ir cells. When significant omnibus effects were detected, Tukey’s HSD post-hoc tests were used.

## Results

### Actigraphy Measures

#### Total Sleep Time

During the white light period, there was a significant interaction between hormone condition and HSP day on cumulative sleep duration (F(12, 141) = 2.702, p = 0.0026, **Figure 1A**). Post-hoc comparisons revealed that on Day 16 (first “late pregnancy” recording), both sustained (p = 0.0284) and withdrawn (p = 0.0002) animals had significantly shorter cumulative sleep durations than oil-treated animals. On Day 17 (second “late pregnancy” recording), sustained animals had significantly shorter sleep durations than oil-treated animals (p = 0.0413, **Figure 1C**). Although withdrawn animals also had decreased total sleep time compared to oil-treated animals, this difference did not reach significance (p = 0.0707, **Figure 1D**). There was no difference across hormone conditions on Day 22 (“postpartum” recording, all p > 0.05, **Figure 1E**). There was also a significant main effect of HSP day on cumulative sleep duration (F(6,141) = 3.117, p = 0.0067). Posthoc comparisons revealed that Day 10 (second “early pregnancy” recording) differed significantly from Day -1 (p = 0.0471).

**Figure 1:**
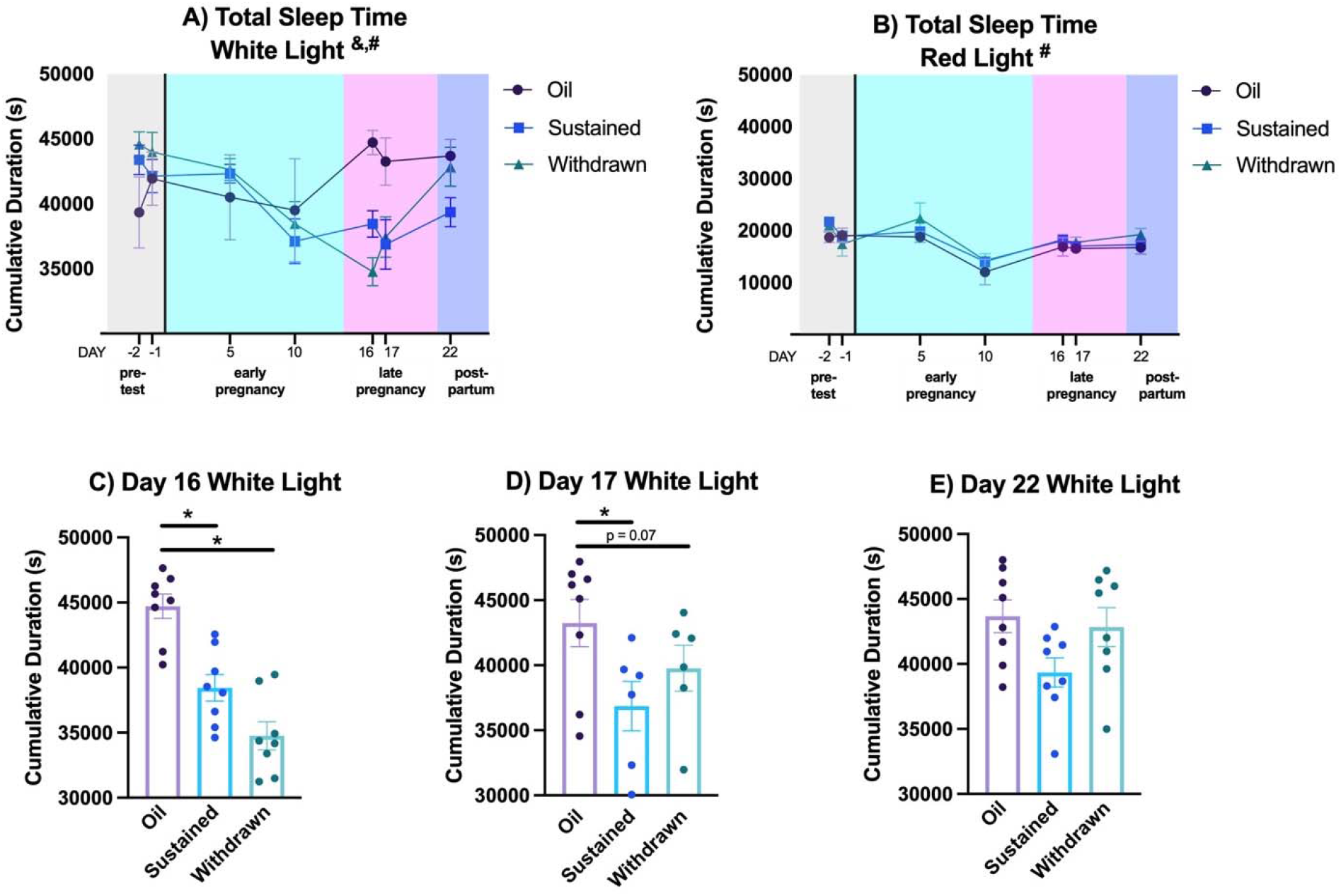
Total Sleep Time. A) During the white light period, there was a significant interaction between hormone condition and HSP day on cumulative sleep duration, as well as a significant main effect of HSP day on cumulative sleep duration. B) During the red light period, there was only a significant main effect of HSP day on cumulative sleep duration. C) Post-hoc comparisons for the White Light period revealed that on Day 16, both sustained and withdrawn animals had significantly shorter cumulative sleep durations than oil-treated animals. D) On Day 17, sustained animals had significantly shorter sleep durations than oil-treated animals. D) Although withdrawn animals also had decreased total sleep time compared to oil-treated animals, this difference did not reach significance. E) There was no difference across hormone conditions on Day 22. Data shown as mean ± SEM. **&** denotes an interaction between hormone condition and HSP Day (p < 0.05); **#** denotes a main effect of HSP Day (p < 0.05); **\*** denotes significant difference across hormone conditions (p < 0.05).

During the red light period, there was a significant main effect of HSP day on cumulative sleep duration (F(6, 141) = 8.479, p < 0.0001, **Figure 1B**). Post-hoc comparisons revealed that Day 10 differed from Day -2 (p < 0.0001), Day -1 (p = 0.0004), Day 10 (p < 0.001), Day 16 (p = 0.0044), Day 17 (p = 0.0423), and Day 22 (p = 0.0040).

#### Sleep Efficiency

During the white light period, there was a significant main effect of HSP day on sleep efficiency (F(6,141) = 2.220, p = 0.0445) and a significant main effect of hormone condition on sleep efficiency (F(2, 141) = 2.995, p = 0.0532, **Figure 2A**). Post-hoc comparisons revealed no significant differences in sleep efficiency across test days. However, during both “late pregnancy” recordings, sustained animals had significantly decreased sleep efficiency compared to oil-treated animals (Day 16 p = 0.0381, **Figure 2C**; Day 17 p = 0.0436, **Figure 2D**). Withdrawn animals did not differ from sustained or oil-treated animals on any test day (all p > 0.05).

**Figure 2:**
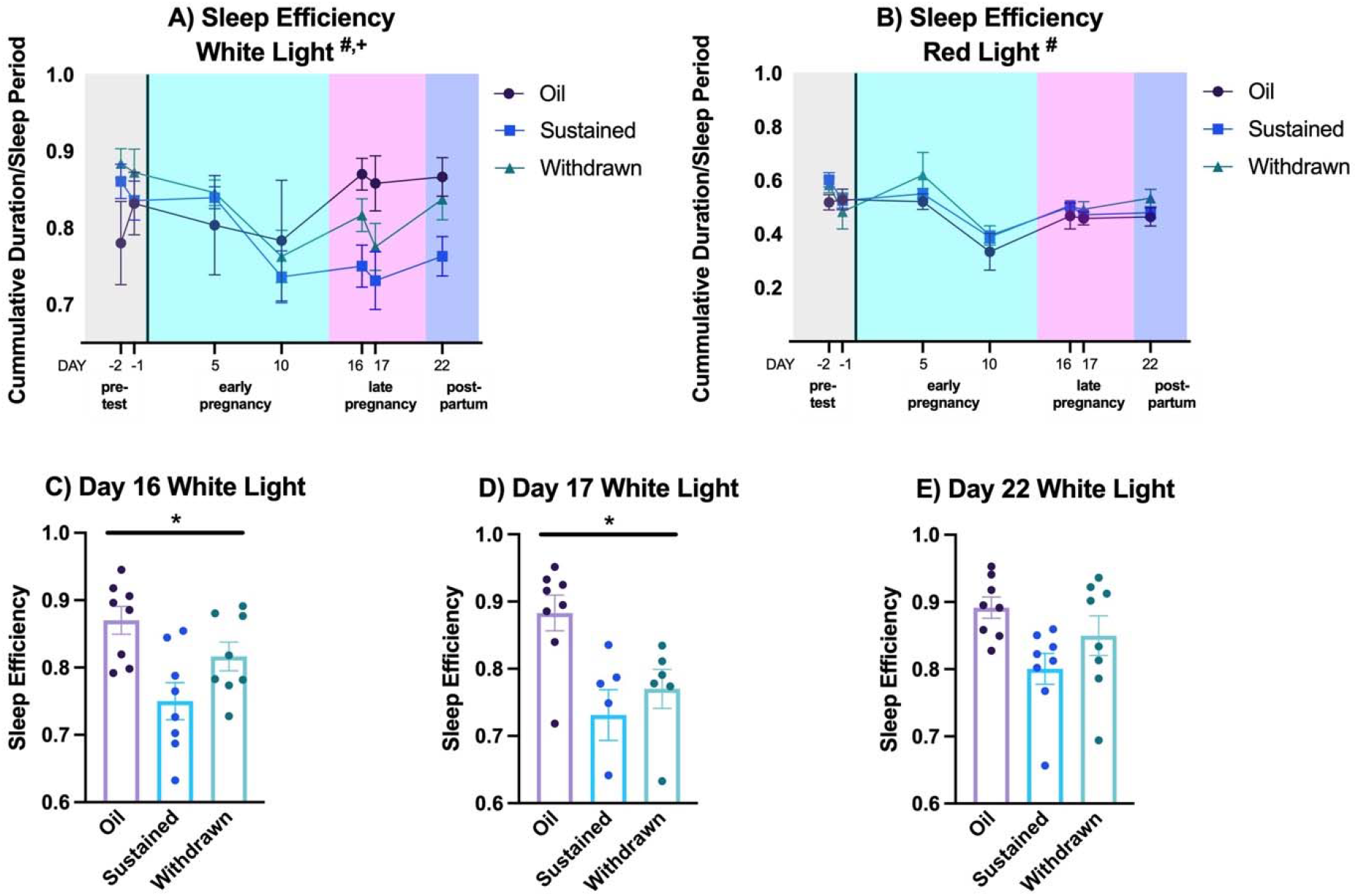
Sleep Efficiency. A) During the white light period, there was a significant main effect of HSP day on sleep efficiency and a significant main effect of hormone condition on sleep efficiency. B) During the red light period, there was only a significant main effect of HSP day on sleep efficiency. Post-hoc comparisons for the White Light period revealed that on C) Day 16 and D) Day 17, sustained animals had significantly decreased sleep efficiency compared to oil-treated animals, whereas withdrawn animals did not differ from sustained or oil-treated animals on either test day. E) There was no difference across hormone conditions on Day 22. Data shown as mean ± SEM. **#** denotes a main effect of HSP Day (p < 0.05); **+** denotes main effect of hormone condition; **\*** denotes significant difference across hormone conditions (p < 0.05).

During the red light period, there was a significant main effect of HSP day on sleep efficiency (F(6,141) = 8.479, p < 0.0001, **Figure 2B**). Post-hoc comparisons revealed that Day 10 differed significantly from Day -1 (p = 0.0004), Day 5 (p < 0.0001), Day 16 (p = 0.0044), Day 17 (p = 0.0423), and Day 22 (p = 0.0040).

#### Sleep Bouts

During the white light period, there was no interaction between hormone condition and HSP day on the number of sleep bouts (F(12,141) = 1.104, p = 0.3612, **Figure 3A**). Likewise there were no main effects of either hormone condition (F(2, 141) = 1.802, p = 0.1687) or HSP day (F(6, 141) = 1.128, p = 0.3491) on the number of sleep bouts.

**Figure 3:**
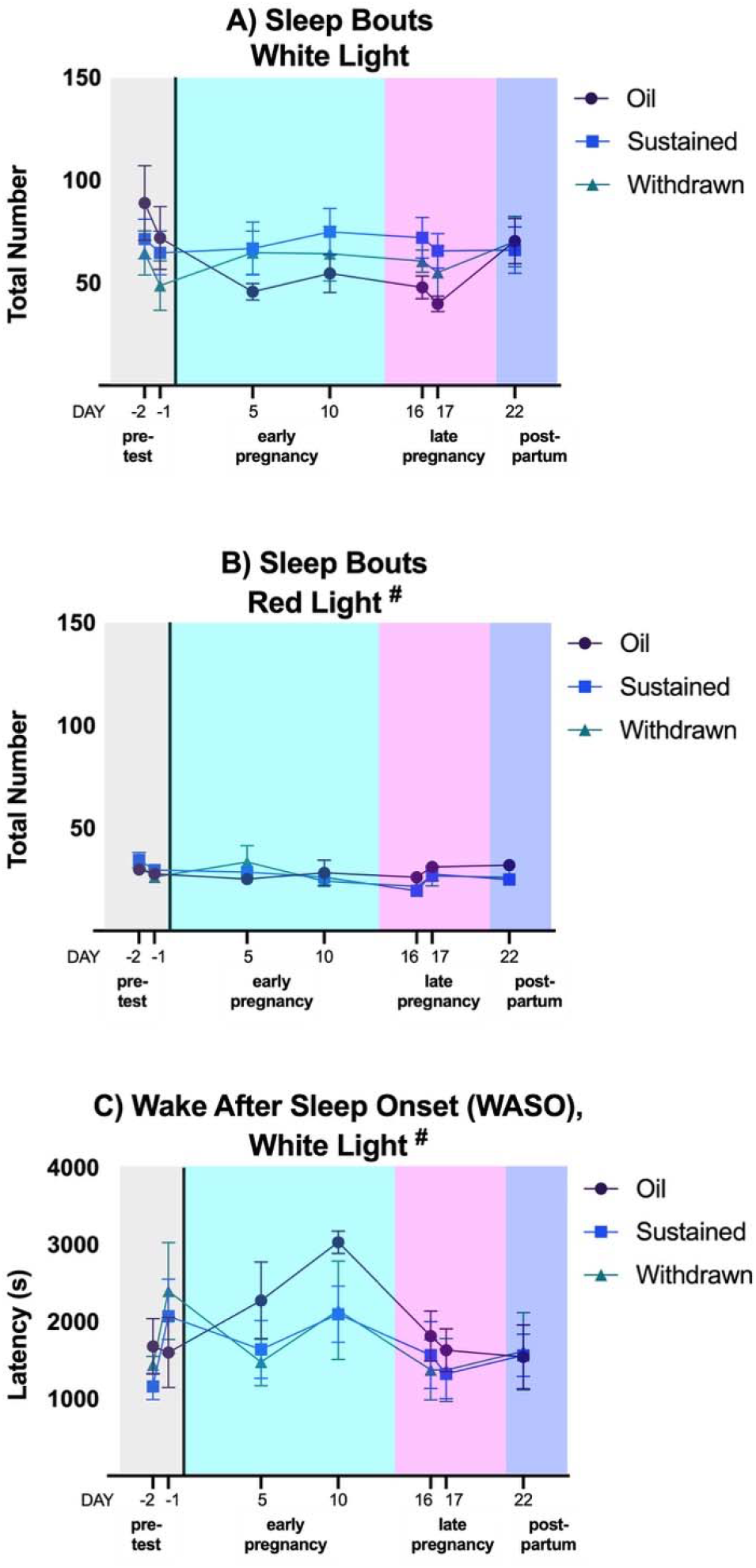
Sleep Bouts and WASO. A) During the white light period, there was no interaction between hormone condition and HSP day on the number of sleep bouts. Likewise there were no main effects of either hormone condition or HSP day on the number of sleep bouts. B) During the red light period, there was a main effect of HSP day on the number of sleep bouts. C) There was a main effect of HSP phase on WASO. However, post-hoc comparisons revealed no significant differences in WASO across test days. **#** denotes a main effect of HSP Day (p < 0.05).

During the red light period, there was a main effect of HSP day on the number of sleep bouts (F(6, 141) = 2.800, p = 0.0132, **Figure 3B**). Post-hoc comparisons revealed that Day -2 differed from Day 16 (p = 0.0030).

#### WASO

During the white light period, there was a main effect of HSP phase on WASO (F(6, 141) = 2.267, p = 0.0404, **Figure 3C**). However, post-hoc comparisons revealed no significant differences in WASO across test days. WASO was not calculated for the red light phase.

#### Distance Moved

During the white light phase, there was a significant interaction between hormone condition and HSP phase on distance moved (F(12, 140) = 2.699, p = 0.0026, as well as main effects of both hormone condition (F(2, 140) = 16.75, p < 0.0001) and HSP phase (F(6, 140) = 4.414, p = 0.0004) (**Figure 4A)**. Post-hoc comparisons revealed that on Day 5 (first “early pregnancy” recording), sustained animals moved significantly farther than oil treated animals (p = 0.0035, **Figure 4C**). On day 16 (first “late pregnancy” recording), both sustained (p < 0.0001) and withdrawn (p = 0.0045) animals moved farther than oil-treated animals, whereas sustained and withdrawn animals did not differ from each other (p = 0.2121) (**Figure 4D**). On day 17 (second “late pregnancy” recording), sustained animals moved significantly farther than oil treated animals (p = 0.0023), but withdrawn animals did not differ from oil-treated animals (p = 0.1139) or sustained animals (p = 0.3807) (**Figure 4E**). Finally on Day 22 (“postpartum” recording), sustained animals moved significantly farther than oil-treated animals (p = 0.0021), whereas withdrawn animals did not differ significantly from oil-treated (p = 0.1106) or sustained animals (p = 0.3757) (**Figure 4F**).

**Figure 4:**
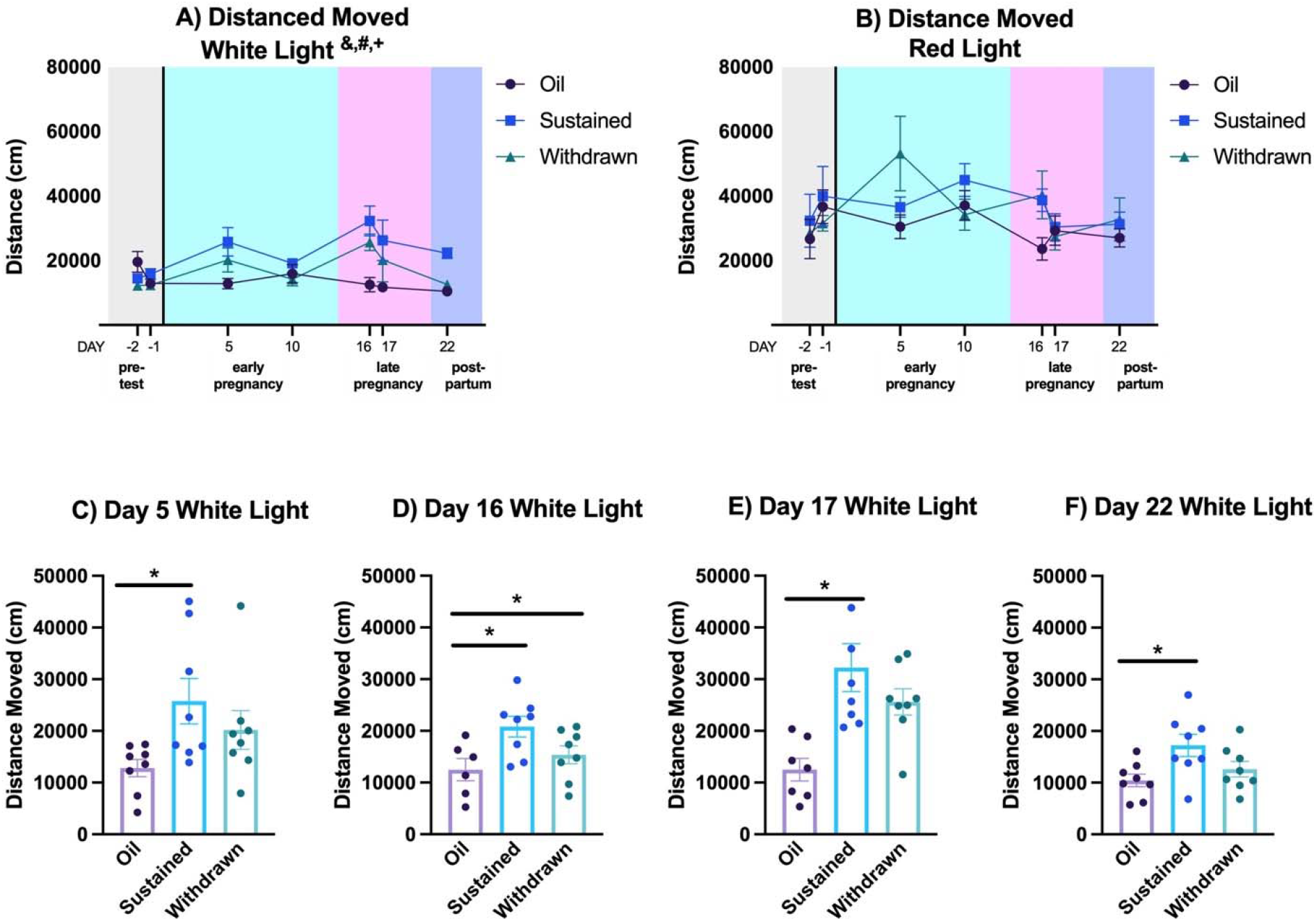
Distance Moved. A) During the white light phase, there was a significant interaction between hormone condition and HSP phase on distance moved, as well as main effects of both hormone condition and HSP phase. B) During the red light period, there was no interaction between hormone condition and HSP phase, although there were trends towards significant main effects of both hormone condition and HSP phase on distance moved. Post-hoc comparisons for the White Light period revealed that C) on Day 5, withdrawn animals moved significantly farther than oil-treated animals. D) On day 16, both sustained and withdrawn animals moved farther than oil-treated animals, whereas sustained and withdrawn animals did not differ from each other. E) On day 17, sustained animals moved significantly farther than oil treated animals, but withdrawn animals did not differ from oil-treated animals or sustained animals. F) On Day 22, sustained animals moved significantly farther than oil-treated animals, whereas withdrawn animals did not differ significantly from oil-treated or sustained animals. Data shown as mean ± SEM. **&** denotes an interaction between hormone condition and HSP Day (p < 0.05); **#** denotes a main effect of HSP Day (p < 0.05); **+** denotes main effect of hormone condition; **\*** denotes significant difference across hormone conditions (p < 0.05).

During the red light period, there was no interaction between hormone condition and HSP phase (F(12, 140) = 1.114, p = 0.3537, **Figure 4B**), although there were trends towards significant main effects of both hormone condition (F(2, 140) = 2.603, p = 0.0777) and HSP phase (F(2, 140) = 1.966, p = 0.0746) on distance moved. Post-hoc comparisons revealed that on Day 5, withdrawn animals moved significantly farther than oil-treated animals (p = 0.0137). There were no significant differences in distance moved across test days (all p > 0.05).

### Brain Measures

#### Ventral Lateral Preoptic Area

In the rostral VLPO, there was a significant effect of hormone condition on the number of cFos-ir cells (F(2, 21) = 3.729, p = 0.0411, **Figure 5A**). Post hoc tests revealed that sustained females had significantly more cFos-ir cells than oil-treated animals (p = 0.0437). In contrast, withdrawn animals did not differ from either sustained (p = 0.6827) or oil-treated (p = 0.1757) females. Similarly, in the caudal VLPO, there was a significant effect of hormone condition on the number of cFos-ir cells (F(2, 21) = 4.428, p = 0.0249, **Figure 5B**). Post hoc tests revealed that sustained females had significantly more cFos-ir cells than oil-treated animals (p = 0.0266). In contrast, withdrawn animals did not differ from either sustained (p = 0.6315) or oil-treated (p = 0.1325) females.

**Figure 5:**
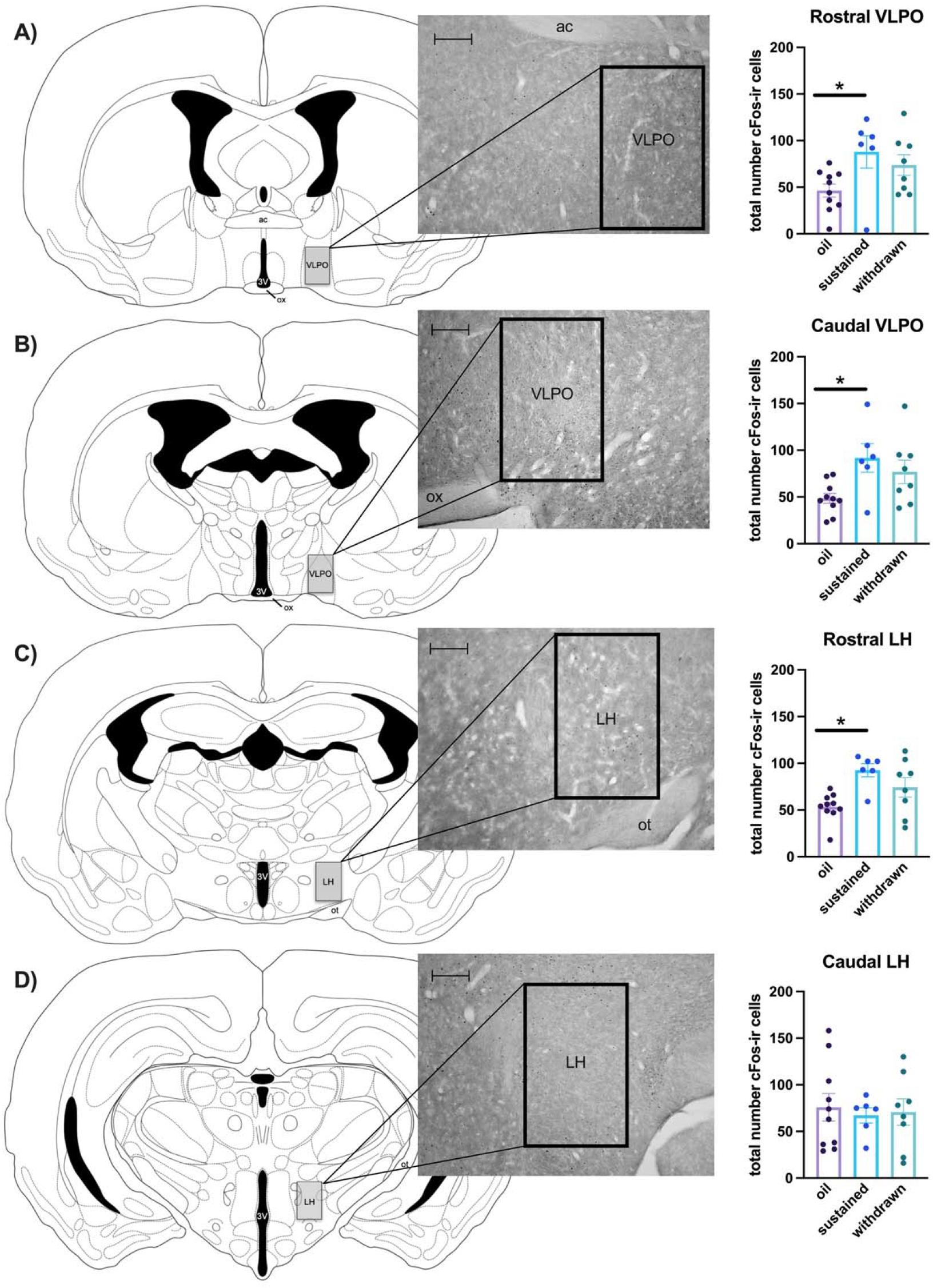
cFos in the VLPO and LH. Counting domains and representative photomicrographs shown for each region of interest (scale bars = 100 μm). A) In the rostral VLPO, there was a significant effect of hormone condition on the number of cFos-ir cells. Post hoc tests revealed that sustained females had significantly more cFos-ir cells than oil-treated animals. In contrast, withdrawn animals did not differ from either sustained or oil-treated females. B) Similarly, in the caudal VLPO, there was a significant effect of hormone condition on the number of cFos-ir cells. Post hoc tests revealed that sustained females had significantly more cFos-ir cells than oil-treated animals. In contrast, withdrawn animals did not differ from either sustained or oil-treated females. C) In the rostral LH, there was a significant effect of hormone condition on the number of cFos-ir cells. Post hoc tests revealed that sustained females had significantly more cFos-ir cells than oil-treated females. In contrast, withdrawn animals did not differ from either sustained or oil-treated females. D) In the caudal LH, there was no significant effect of hormone condition on cFos-ir cells. Data shown as mean ± SEM, **\***denotes significant difference across hormone conditions (p < 0.05).

#### Lateral Hypothalamus

In the rostral LH, there was a significant effect of hormone condition on the number of cFos-ir cells (F(2, 21) = 6.441, p = 0.0066, **Figure 5C**). Post hoc tests revealed that sustained females had significantly more cFos-ir cells than oil-treated females (p = 0.0054). In contrast, withdrawn animals did not differ from either sustained (p = 0.2786) or oil-treated (p = 0.1256) females. In the caudal LH, there was no significant effect of hormone condition on cFos-ir cells (F(2, 21) = 0.09834, p = 0.9068, **Figure 5D**).

## Discussion

Here we demonstrate for the first time that increased estradiol during a simulated pregnancy decreases total sleep time and sleep efficiency while increasing home cage activity in female Syrian hamsters. These data map well onto clinical data showing that sleep is disrupted during late pregnancy^6–8^. Importantly, the HSP model used in this study allows for the dissociation of estradiol’s direct impact on sleep, presumably via action in the brain, from the impact of other hormonal changes and physical/environmental factors associated with pregnancy and parental care. Using this model, we found that high levels of estradiol increase the expression of cFos, an indirect marker of neuronal activation, in the VLPO and LH compared to ovariectomized females. In contrast, animals who were withdrawn from estradiol during the simulated postpartum phase did not differ from oil-treated controls in either behavior or brain measures. Together, these data suggest that increased levels of estradiol during pregnancy act in estrogen-sensitive hypothalamic sleep nuclei to influence sleep during the peripartum period.

Syrian hamsters are a well-studied model in sleep and chronobiology research due to their robust and predictable circadian rhythms^56^. They are also an excellent choice for behavioral actigraphy studies like the present one because, unlike other common rodent models, they prefer to be single-housed^40,41^, facilitating accurate video tracking in the home cage environment. While actigraphy lacks the specificity of other techniques such as electroencephalogram (EEG) and electromyogram (EMG), it is a non-invasive and relatively low-cost option that has been shown to track basic sleep parameters with 90-95% agreement with EEG/EMG recordings^57^. Nonetheless, there are limitations to this method. For example, in the present study we found that high levels of estradiol decreased total sleep time and sleep efficiency during “late pregnancy,” but we were unable to detect a behavioral mechanism for these decreases; that is, although hormone-treated animals appeared to have more sleep bouts and shorter latencies to wake (WASO) than oil-treated animals during late pregnancy, these differences did not reach statistical significance. It is likely that a more sensitive method, such as EEG/EMG, may be required to detect differences in measures like sleep bouts or WASO that would contribute to an overall decrease in sleep time.

Our finding that high levels of estradiol during “late pregnancy” decreased actigraphy measures of sleep is supported by data from pregnant people in which sleep disruptions are worst during late pregnancy, when estrogen levels are highest^6–8^. Our data also corroborate data from across the rodent estrous cycle^26,58–60^, in which sleep is inversely related to estrogen levels. Interestingly, total sleep time is generally *increased* during pregnancy in rodents^60–62^, with a particular increase in non-rapid eye movement (nREM) sleep during late pregnancy^61,62^. This increase is thought to result from the increased metabolic load of pregnancy, which may necessitate more sleep. However, late pregnancy in rodents is also characterized by frequent switches between wakefulness and nREM sleep^62,63^, suggesting sleep fragmentation. In this context, our data support the hypothesis that high levels of estrogens during late pregnancy suppress sleep, but this suppression may present differently in the absence of other physiological factors associated with pregnancy. These hormone-mediated sleep disruptions may be adaptive, particularly in primiparous individuals, in preparing for postpartum sleep disruptions associated with parental care.

In the present study, the effects of hormone treatment on sleep parameters were found primarily during the white light phase. This differs from previous findings in ovariectomized rats, in which treatment with estradiol and/or progesterone reduce nREM and REM sleep and increase brief awakenings during the dark phase only^28^. This discrepancy may be explained by differing levels of hormones; the silastic capsules in the previous study generate hormone levels typical of diestrus and proestrus, whereas the daily hormone injections in the current study generate much higher levels of estradiol and progesterone that are meant to induce maternal behavior in nulliparous females^46^. In addition to differing hormone levels, different light:dark cycles (14:10 in the current study versus 12:12 in the previous study) may influence the study outcomes. Simply put, the effects of hormone treatment on less-sensitive actigraphy measures may have been easier to detect during the longer light phase than in the shorter dark phase. It is noteworthy, however, that the impact of estradiol treatment on circadian rhythms does not differ between Syrian hamsters housed at 12:12 versus 14:10 light:dark cycles^64^. Our results may therefore reflect a species difference in the impact of estradiol treatment on sleep behavior between seasonal vs. non-seasonal breeders.

It is well established in non-pregnant rodents that heightened levels of estradiol, either via endogenous fluctuations^60,65^ or exogenous administration^66–71^, increase locomotor activity. These studies have most commonly used voluntary wheel running to assay activity, but estradiol has also been reported to enhance activity in the home cage and open field^72^. Progesterone, on the other hand, has no effect on voluntary activity levels in Syrian hamsters; however when given in combination with estradiol, it antagonizes estradiol-induced increases in activity^66^, supporting our finding that increased locomotion in “early pregnancy” (estradiol plus progesterone) is lower than in “late pregnancy” (estradiol alone). During pregnancy, voluntary wheel running is dramatically lowered in rats in mice ^73,74^. In these species, estradiol’s enhancing effect on voluntary wheel running depends on activation of estrogen receptors in the medial preoptic area (MPOA)^67,68^, whereas pregnancy-induced reductions in voluntary activity are mediated by prolactin action in the MPOA^75^. Taken together, this suggests that during pregnancy, prolactin must overcome the pro-ambulatory actions of estrogens in the MPOA in order to suppress voluntary activity. Unlike in rats and mice, however, voluntary wheel running remains unaffected throughout most of pregnancy in Syrian hamsters, decreasing only in the last three days of pregnancy^76^. While voluntary wheel running and sleep are typically inversely related, this relationship breaks down in hamsters during pregnancy^60^. Given this species difference, as well as the lack of prolactin surge in the HSP model, it is not surprising that we found that estradiol treatment, alone or in combination with progesterone, increased distance traveled in the home cage. These data provide unique insight into the impact of peripartum hormone fluctuations on voluntary activity, and suggest that Syrian hamsters may be an advantageous animal model in which to investigate the pathophysiology of RLS, which is characterized by an urge to move the legs that is worsened during the night or while resting^77^, during pregnancy.

While the activity-promoting effects of estradiol are likely localized in the MPOA, the effects of estradiol on sleep duration and efficiency may be housed elsewhere in the hypothalamus. We found that animals that continued to receive high levels of estradiol through the simulated postpartum period had more cFos-ir cells in the caudal LH, rostral VLPO, and caudal VLPO compared to oil-treated females. It is straightforward to understand how decreased sleep would correlate with increased activation of the LH, an arousal-promoting nucleus^78^. Wake-promoting orexin/hypocretin neurons in the LH are sensitive to fluctuations in ovarian hormones^79^. Less straight-forward, however, is the finding that estradiol simultaneously increased activation of the VLPO, a sleep-promoting nucleus^78^. In fact, previous studies in rats have shown that estradiol treatment, albeit at lower levels, decreases cFos expression^35^ and other molecular markers of neuronal activation^36^ in the VLPO. It is noteworthy, however, that in ovariectomized rats, sleep deprivation increases cFos expression in both the LH and VLPO, and estradiol treatment does not impact cFos expression in the VLPO in sleep deprived animals^39^. It is therefore possible that in the current study, changes in cFos expression in the VLPO are a consequence of sleep deprivation rather than a driving force behind it. In support of this idea, animals that were withdrawn from estradiol to simulate the rapid decline in estrogens following parturition had intermediate levels of both sleep disruption and cFos expression compared to sustained and oil-treated females. The impact of pregnancy-like levels of estradiol on sleep may be mediated by other estrogen-sensitive sleep regulatory nuclei that project to the VLPO, such as the dorsal raphe^80^. Indeed, the DRN has previously been shown to be plastic in response to HSP in Syrian hamsters^54^. In the future, more comprehensive brain region sampling, as well as identifying the neurochemical phenotype and connection patterns of cFos-ir cells, is needed to fully elucidate the impact of peripartum estradiol fluctuations on sleep-associated neural circuitry and behavioral outcomes. Nonetheless, the present results suggest that high levels of estradiol during late pregnancy act in the LH and, and possibly indirectly in the VLPO, to suppress sleep during pregnancy. Ultimately, these data help to differentiate the impact hormonal versus other physiological, anatomical, and/or environmental factors on peripartum sleep disruptions.

## Acknowledgements

This work was funded in part by a Teaching with Technology Grant to LEB from Haverford College. The authors wish to thank Jonathan Dale at Noldus Technologies for his help setting up Ethovision to track actigraphy measures.

